# Postdoctoral Scholar Recruitment and Hiring Practices in STEM: A Pilot Study

**DOI:** 10.1101/2024.02.12.580018

**Authors:** Meagan Heirwegh, Douglas C. Rees, Lindsey Malcom-Piqueux

## Abstract

Despite the importance of the postdoctoral position in the training of scientists for independent research careers, few studies have addressed recruiting and hiring of postdocs. We conducted a pilot study on postdoctoral hiring in the Division of Chemistry and Chemical Engineering at the California Institute of Technology to serve as a starting point to better understand postdoctoral recruiting and hiring processes. From this survey of both postdocs and faculty, together with the available literature, the picture emerges that the postdoc hiring process is more decentralized than either faculty hiring or graduate admissions. Postdoc positions are often filled through a passive process where the initial expression of interest from a prospective postdoc is through a “cold-call” contact to a prospective advisor. Individual faculty members are often responsible for developing and implementing their own outreach and recruitment plans and deciding who to hire into a postdoc position. The overall opacity of the processes and practices by which postdocs are identified, recruited, and hired make it difficult to pinpoint where interventions could be effective to ensure equitable hiring practices. Implementation of such practices is critical to training a diverse postdoc population and subsequently of the future STEM faculty recruited from this group.

## Introduction

The postdoctoral (postdoc) appointment represents an important step on the path to an independent career in research (National Academy of Sciences (2014)), which can include securing a STEM faculty position in a research university. Indeed, an estimated 50-70% of scholars intend to go into a tenure track position at the start of their postdoc, with ∼10-15% actually securing a life or physical sciences or engineering tenure track position within 5 years of completing their doctorate (National Academy of Sciences (2014), Sauermann and Roach (2016), Andalib, Ghaffarzadegan et al. (2018), Grinstein and Treister (2018), Denton et al. (2022)). Consequently, finding and completing a productive postdoctoral appointment are critical steps on the pathway to the STEM professoriate at research universities. As a corollary, given role of the postdoctoral positions in the preparation of faculty, equitable postdoc hiring practices are critical to diversifying the STEM faculty (see Matias et al, 2022, Patt et al, 2022).

While multiple studies have focused on the postdoc experience, duration and career path (National Academy of Sciences (2000), National Academy of Sciences (2014), Eaton, Saunders, et al. (2019), Lambert, Wells et al. (2020), König (2022)), few studies have examined the recruitment and hiring practices used to get scholars into postdoctoral positions. A significant challenge to these studies is the inadequate and incomplete data collection on the postdoctoral population (National Academy of Sciences (2014)) including information on demographics, career aspirations and career outcomes, as well as how hiring decisions are made. This dearth of information reflects the highly decentralized nature of the postdoctoral position, with funding and mentoring often the responsibility of individual faculty. As noted by Huynh and Shauman (2021), “universities typically provide little to any guidance or oversight of postdoctoral hiring. The direct employers are individual researchers who fund the postdoctoral positions from grants obtained externally” reflecting “the notable absence of explicit and consistent policy about, guidelines for, and oversight of postdoctoral hiring”. This situation stands in stark contrast to the more collective responsibility of, and oversight by, departments, colleges, universities for the admission of undergraduate and graduate students, and for the hiring of faculty (Liera and Ching, 2020, National Academies of Sciences, Engineering, and Medicine, 2023, National Association of College Admissions Counseling, 2023, Posselt (2016, 2020), Sensoy and DiAngelo (2017)).

The appointment of a postdoc by a faculty member involves multiple considerations and steps by both the prospective postdoc and the prospective faculty advisor. These can include whether to advertise a position, whom to interview and whom to hire on the part of the prospective advisor, and where to apply and whether to accept an offer on the part of the prospective postdoc. A short summary of postdoctoral hiring, based on focus group discussions with several hundred postdocs, faculty, administrators, etc. was presented by the National Academy of Sciences (2000). Two approaches to the hiring process were highlighted (i) personal contacts (advisors, friends, contacts from meeting) and (ii) directly approaching potential advisers with their qualifications and research interests. Significantly, responding to posted positions was not seen as an effective strategy for finding a postdoc. A survey of the MIT postdoctoral community (MIT Postdoctoral Association, (2020)) provided quantitative support for these observations, with 44% of postdocs obtaining their position through a personal or work connection, while 43% obtained their position through a cold call. Only 11% of the postdocs found their position by responding to a job position.

The recent report by Huynh and Shauman (2023) represents an important advance in understanding the hiring process for postdoctoral scholars. This study analyzed the relationship between gender, race-ethnicity and hiring of postdocs in a state university system. A number of important findings were reported, including that while women from underrepresented racial/ethnic groups are more likely to be invited to interview for a postdoc position, they are also less likely to receive an offer when compared to all interviewed candidates. Of note, this study further established that there are significant variations by gender and race-ethnicity in who applies for postdoctoral position, and this variation, not surprisingly, impacts who is hired into these positions. Although employed by only a minority of candidates (6.2%) in this analysis, candidates who learned about a postdoctoral position through their personal networks experienced a significant enhancement in securing that position.

Though some studies have explored the extent to which gender biases and racial biases shape individual faculty members’ assessments of postdoctoral candidates in ways that may hurt some candidates’ chances of being hired (e.g., Eaton, Saunders, et al. (2020)), it is not yet clear how the recruitment process and hiring practices might exacerbate inequity by further disadvantaging prospective postdocs from historically excluded and minoritized groups. Similarly, little has been published about how postdoctoral candidates experience their postdoc job search, including whether these experiences diverge for candidates from historically excluded and minoritized populations.

### A Pilot Study

In view of both the importance of, and the lack of information about, the postdoctoral hiring process, we conducted a pilot study to serve as a starting point to better understand this process and to identify some of the critical aspects. Faculty and postdocs appointed in the Division of Chemistry and Chemical Engineering (CCE) at the California Institute of Technology (Caltech) were invited to participate in anonymous, voluntary online surveys. All participants completed informed consent. This study was reviewed by the Caltech Institutional Review Board (protocol #IR22-1240) and was determined to be exempt from full review. CCE was selected as a smaller, self-contained test case for the campus as a whole. The division has a large postdoc population working in wide range of subject areas that often overlaps with work in other divisions at Caltech.

In general, postdoc hiring at Caltech is not centrally managed by the institute. There is not a central postdoc job board for all positions nor does Human Resources manage or coordinate the search for candidate. Very few postdoc positions are advertised centrally. Institutional structures, such as the Caltech Postdoc Office, offer indications that postdocs are hired at Caltech but do not explicitly advertise positions or detail how one would go about securing a position. Most postdoc hiring is done at the individual faculty level, at the discretion of each of these faculty members. These positions are often paid through research grants within a group or by external postdoc funding. Faculty members are able to decide how and when to go about finding postdocs with minimal institutional oversight or guidance. Some positions, such as prize postdoctoral fellowships within one division or department, are overseen by a committee. Often these positions are funded by a prize, endowment or central departmental funds. There is no standard, universal process used across Caltech when recruiting and hiring postdocs. Across peer institutions, Caltech is not unique in this approach to postdoc hiring (see Huynh and Shauman (2021)). Hiring at the discretion of a single faculty member is ubiquitous at these institutions.

The surveys used were designed to examine both sides of the postdoc hiring process, from the faculty and postdoc perspectives. Separate survey instruments were created for the two populations (Appendix 1). Each instrument consisted of closed and open-ended items relating to academic and career backgrounds, self-identified sex, methods used to find current and previous postdoc positions, methods used to identify and recruit postdocs (faculty only) and perceptions of effective hiring practices. Participation from both faculty and postdoc was relatively high with 77% (34/44) of faculty members and 61% (65/106) of postdocs choosing to participate. Note that not all questions were required and not all participants chose to answer all questions, so the number of responses may differ between questions. A summary of the survey results is provided in Appendix 2; through this paper, responses to survey questions in Appendix 2 will be represented as FN or PN, where N denotes the question number on the faculty (F) or postdoc (P) survey, respectively,

### Limitations

This survey reflects the experiences of faculty and postdocs in one division (CCE) at one university (Caltech). As Caltech is a small institution, the sample size is accordingly small and likely not fully representative of the range of experiences in postdoc hiring. Critically, this survey was also only able to reach postdocs who had successfully navigated the hiring process and were employed. There is no way, in this survey, to capture the experiences of those who were unsuccessful or self-selected out of the process. Thus, the responses discussed here do not fully paint a picture of the entire hiring process.

### Faculty Demographics

Of the 34 faculty participants, 21% (7/34; F1) had earned their PhD (i.e., had an academic age) in the past 9 years while 32% (11/34; F1) had earned their PhD 40 or more years ago, suggesting that a wide range of faculty experience and seniority had been captured. Reflecting the importance of the postdoc position, 82% (28/34; F3) of the faculty held a postdoc prior to their first faculty appointment, while 93% (26/28; F8) of the faculty report having postdocs in their research group at present and/or previously.

### Postdoc Demographics

The majority of Caltech postdoc respondents did not complete their undergraduate education in the United States (69%, (45/65; P1) whereas half (51%, (33/65; P2)) completed graduate school here. For 75% (49/65; P3) of postdocs, the position they currently held at Caltech was their first postdoc appointment. Of the 16 postdocs who had held a previous postdoc appointment, 9 were with the research group in which they had completed their PhD (P4), suggesting that their Caltech postdoc the first time they had to seek out a postdoc position. Most (59%, (37/63; P8)) postdocs had been at Caltech as a postdoc for less than 2 years.

## Results and Discussion

From this pilot project, we found that postdoc positions are most likely filled through an essentially passive process where a prospective postdoc initiates contact with a prospective advisor. 69% (18/26; F13) of the faculty reported this approach resulted in the most hires (Figure 1), which is consistent with the summary of this process reported in the National Academy of Sciences (2000) assessment. Hence, the major approach resulting in the greatest number of hires is a passive method that does not involve the prospective advisor posting any notices of potential postdoc opportunity. Recommendations from collaborators or other peers, another passive method, was the second most reported method (15%, (4/26; F13)). Active methods – such as including a statement on a research group website, advertising on social media, or other forms of advertising available postdoc positions – accounted for the remaining responses (15%, (4/26; F13)).

**Figure 1.**
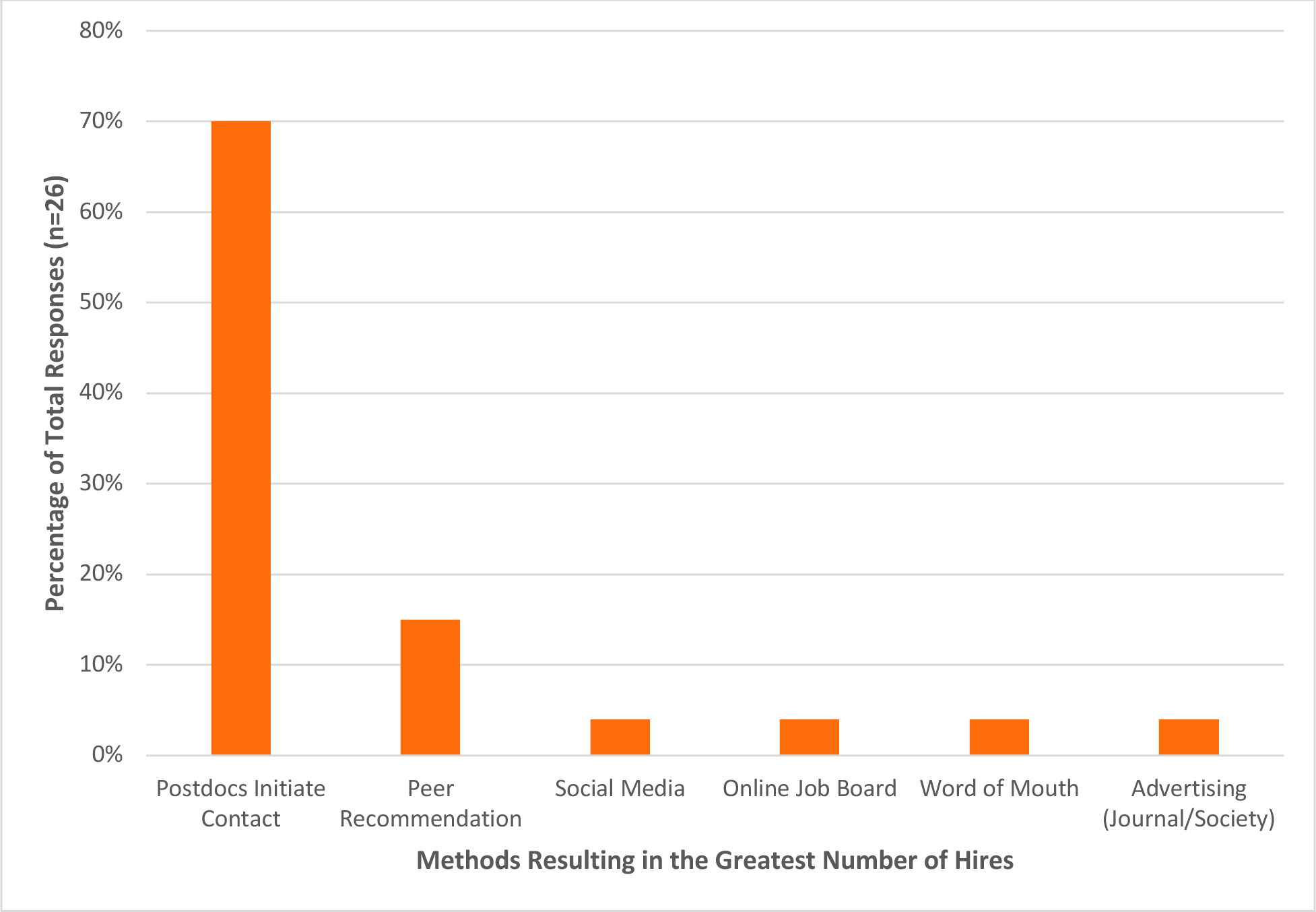
Caltech faculty responses when asked which recruitment method has resulted in the greatest number of postdocs hires in the past 15 years (F13). Respondents could select only one response.

The process by which Caltech postdocs found their current position mirrors the faculty experience, although with some important differences (Figure 2; P9). The most common mechanisms by which current postdocs found their positions were a referral from their PhD advisor/mentor (29%, (18/63), a cold-call application to a prospective advisor (19%, (12/63)), applying to a posted opportunity (14%, 9/63), and previous research-based contact with a prospective advisor (13% (8/63)). While the first two reflect the passive approaches from advisors noted in the faculty survey, the latter two require a more active participation in the recruitment process by the future postdoc advisor. These experiences also reinforce that the PhD advisor can play an important role in helping their advisee secure a postdoc position, either through referrals or research collaborations. Although 65% (41/63; P13) report that their PhD advisor did not reach out to their network to help them find a position, when the advisor did reach out a potential position resulted 85% (19/22; P14) of the time. A direct referral from a peer faculty member at another institution may help the prospective postdoc stand out from others looking for a postdoc. The MIT postdoc survey (2020) found a similar trend. 25.3% of respondents were introduced or recommended to their postdoc advisor via a connection. Though they did not specifically ask about PhD supervisors, they would fall into this category. A pre-established connection, whether their own or through a member of their network, can be extremely helpful when finding a postdoc position.

**Figure 2.**
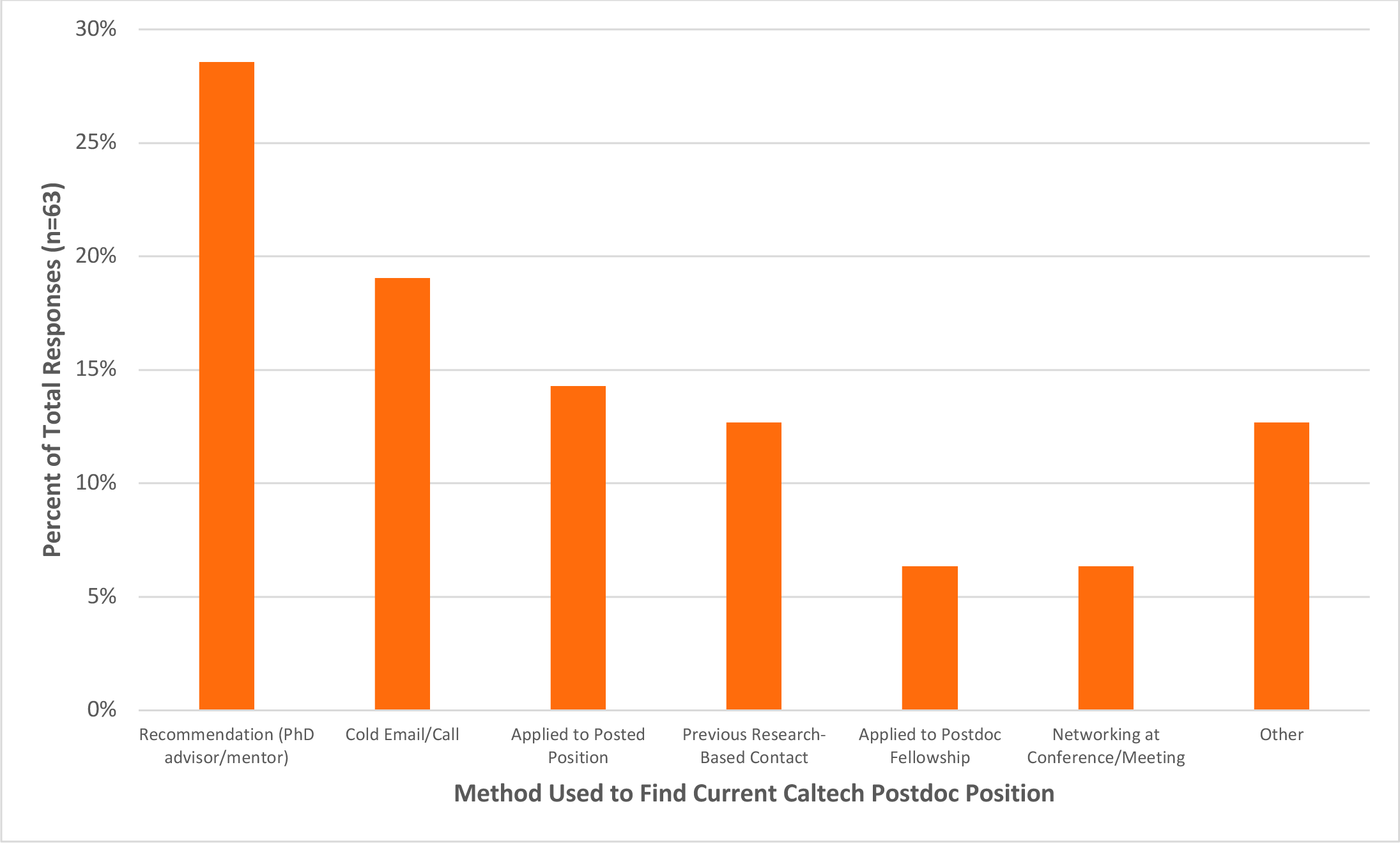
Methods used by Caltech CCE postdocs to find their current postdoc position at Caltech (P9). Respondents could select only one response.

As a reference, faculty were asked how they found their own postdoc position when they were at that career stage (Table 1). 41% (13/32; F6&7) of faculty found their own postdoc positions by either a cold phone call or email to a scholar in the field. This contrasts with the current Caltech postdocs, where the most common way that they found their positions was from a referral from their PhD advisor/mentor (29%, 18/63; P9), while applying to a posted opportunity was second most common method used to find a position (19%, 12/63; P9)). Notably, no faculty member who responded to this survey applied for a posted postdoc position. It should be noted that faculty with an academic age of 25 or more years likely did not have access to a significant number of online postings.

**Table 1.**
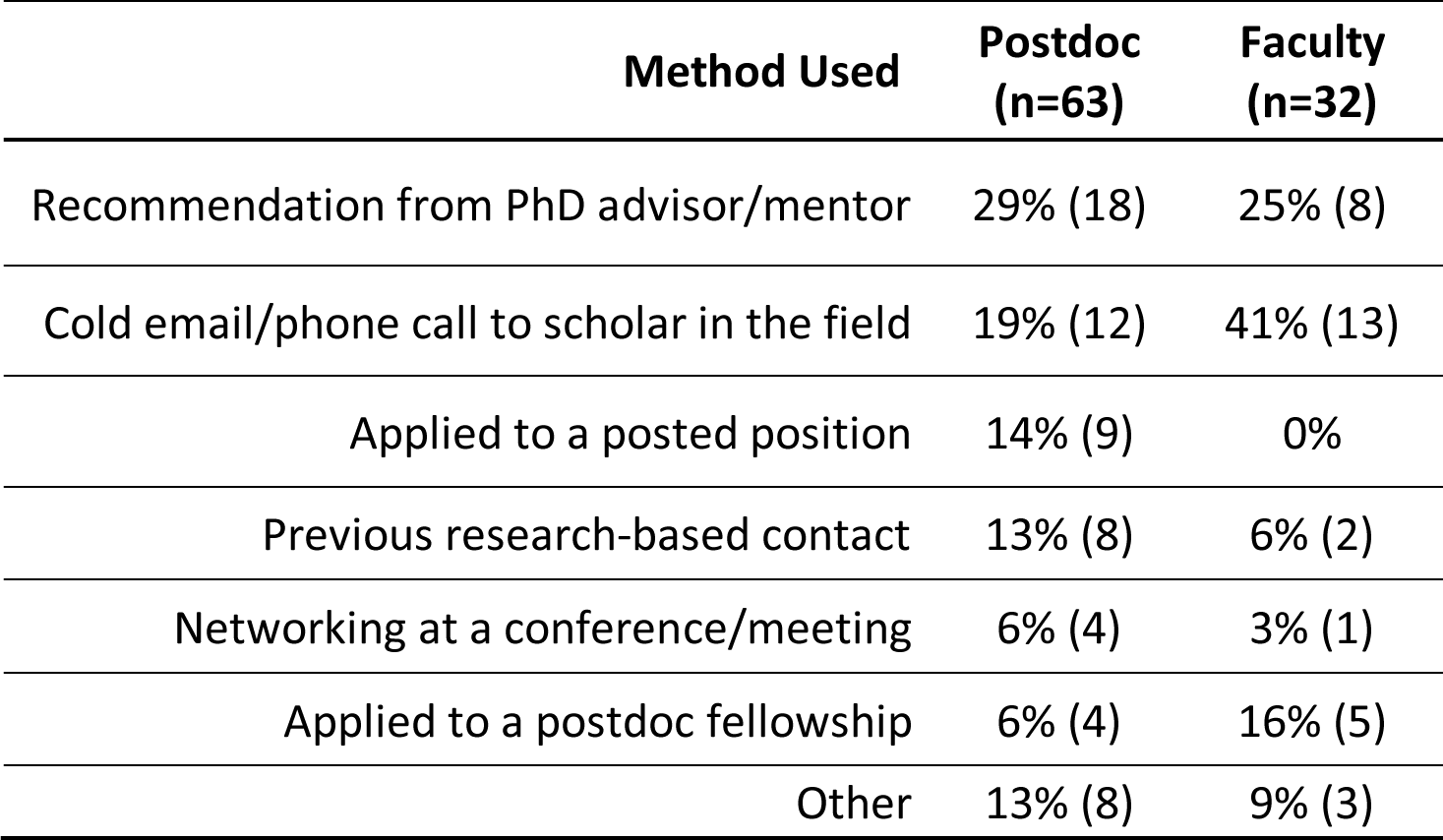
Comparison of the methods used by current CCE postdocs (P9) and faculty (when they were postdocs; F6&7) to find their postdoc positions. Respondents could select multiple responses, if different methods were used to find multiple postdoc positions. Percentages are reported as the percent of total responses.

Faculty were asked three related questions about the methods they use to find postdocs for their research groups, with the results summarized in Table 2:

i. all methods currently used to find postdocs (F11; n = 82)
ii. all methods used to hire the postdocs currently in the research group (F15; n = 44)
iii. methods that have resulted in the greatest number of postdoc hires in the last 15 years (F13; n = 28)

**Table 2.** Faculty response when asked which methods are currently used to find postdocs in the research group. Respondents could select as many responses as needed. (P) and (A) denote passive and active methods, respectively.

| **Method** | **Currently used methods to hire postdocs (F11; n=82)** | **Methods used to hire current postdocs (F15; n=44)** | **Most successful methods to hire postdocs in last 15 years (F13; n=26)** |
| --- | --- | --- | --- |
| Postdocs initiate contact (P) | 26% (21) | 43% (19) | 69% (18) |
| Peer Recommendations (P) | 18% (15) | 18% (8) | 15% (4) |
| Email to Peers (A) | 12% (10) | 7% (3) | 0% |
| Research Group Website (general call) (A) | 11% (9) | 5% (2) | 0 (0%) |
| Research Group Website (specific position) (A) | 7% (6) | 5% (2) |  |
| Social Media (specific position) (A) | 9% (7) | 9% (4) | 4% (1) |
| Social Media (general call) (A) | 6% (5) | 0% (0) |  |
| Advertising (Journal/Society) (A) | 5% (4) | 5% (2) | 4% (1) |
| Online Job Board (A) | 2% (2) | 2% (1) | 4% (1) |
| Other Caltech Website (A) | 1% (1) | 0% (0) | 0% |
| Other | 2% (2) | 7% (3) | 4% (1) |

Respondents were permitted to select more than one method, as needed. Table 2 divides the methods used by faculty to find postdocs into active and passive methods. Passive methods include postdocs initiating contact and recommendation from peers, while all other methods were considered active and involve some mechanism by which faculty announce the availability of postdoctoral positions on a forum. The results were consistent across all three questions that the top two responses for recruiting postdocs are prospective postdocs initiating contact and recommendation from a peer or collaborator. Intriguingly, although these two passive methods are reported by a significant majority of faculty as the most successful method for hiring postdocs (85%, ((18+4=22)/26; right column in Table 2)), active methods constitute a majority of all methods used to recruit postdocs (56%, (46/82); left column in Table 2); 62% faculty (16/26; F10) report having advertised for a specific postdoc position vacancy in the past 15 years. The ineffectiveness of active methods suggests that there is some underlying mechanism (or mechanisms) that works against the successful recruiting of postdocs in this way.

One possible explanation is that there is a difference in utilization of active methods by faculty of different academic ages. For example, social media and websites would not have been available to faculty above a certain academic age at the beginning of their independent careers, and hence they started when active methods were more challenging (requiring mailing letters or posting announcements in a society newsletter). To assess the relevance of this effect, the average number of each type of method used by each academic age group was calculated for the answers to question F11 (Table 3). Unsurprisingly, those faculty who most recently earned their PhDs utilize active recruiting methods most frequently. Not only are these faculty members more likely to use active recruiting methods, but they are also more likely to use multiple methods including social media and posting on their research group websites. While the use of passive methods is relatively similar between the different academic age groups, more variability is evident in the use of active methods. Intriguingly, social media usage was not restricted to younger academic age but was also employed by several faculty in the 30-39 year academic age group as well.

**Table 3.** Average number of active and passive methods used for postdoc recruitment, by academic age. Respondents could select as many methods as were used sometime in the past 15 years (Table 2). Passive methods include 2 options: postdocs initiate contact and recommendations from collaborators, colleagues and other peers. Active methods include mechanisms for posting available positions on some forum as identified in Table 2.

|  | **Academic Age (Years Since Receiving PhD)** |  |  |  |  |
| --- | --- | --- | --- | --- | --- |
|  | **0 - 9** | **10 - 19** | **20 - 29** | **30 - 39** | **40+** |
| **Passive** | 1.3 | 1.5 | 1.7 | 1.3 | 1.5 |
| **Active** | 2.8 | 1.3 | 1 | 2.2 | 0.8 |

At the end of the survey, faculty were asked if there was anything else they would like share about how they find postdocs for their research groups. Two comments addressed experiences in advertising for positions versus interested students initiating contact. One person commented that “[On] a few occasions I have sent out a dear colleague letter, where some specific expertise in my group is desired, but this has not met with great results.”, while a colleague noted “advertisement yields many applications but the best are writing without a posted position”. Though the more active methods may result in a greater number of inquiries about postdoc positions, the sentiment expressed by some faculty members, as well as the responses reported in Table 3 suggests that the most common, passive methods are seen as working for many.

On the other hand, one faculty member noted “It has become increasingly difficult to find postdocs in recent years. I will likely start to advertise on social media and in scientific journals or society”. In what may be a changing postdoc landscape, evidence suggests that postdoc hiring is becoming more difficult (Langin, 2022). Going forward, passive recruitment techniques of relying on interested postdocs to initiate contact or faculty recommendations to appear may not always result in the number of postdocs needed.

Postdocs were asked if they could have pictured themselves as a postdoc at Caltech when they were a graduate student. Their responses were mixed. Several responses mentioned that they had aspired to Caltech because of the reputation and prestige of the institution and the CCE division itself. One postdoc responded, “Working at Caltech as a postdoc was a dream of mine since it is highly ranked in my career field nationally.” Another said “Yes, being a scientist at Caltech has always been a dream, well before I started my PhD. Simple reason being the things you hear about the academic ambience, peer group, exposure to brilliant scientific minds, the resources and technology that are available to name a few.” Others didn’t picture themselves at Caltech because it didn’t seem like an attainable goal, again due to the reputation of the Institute. In a sentiment that was echoed by others, one postdoc responded “No, I didn’t envision being able to land a position at this caliber of institution”. Another postdoc offered more details when they wrote, “I did not originally picture myself ever working at Caltech. The lack of faith can be attributed to my imposter syndrome – particularly, the idea that someone of my background and history could never make it to a prestigious research institute such as Caltech.”

The name-recognition of Caltech appears to be a double-edged sword. For some, the institute and its reputation is enough of a reason to want to apply whereas for others it acts as a source of intimidation. Without clear signals of how to find a postdoc from faculty members, the reputation itself may be enough to deter potential applicants especially given the prevalence of passive recruitment techniques.

### Conclusions & Future Implications

Overall, the picture emerges that hiring a postdoc is more diffuse and decentralized than either faculty hiring or graduate admissions. Instead of relying on search or admissions committees, individual faculty members are often responsible for developing and implementing their own outreach and recruitment plans and use their own discretion when deciding whom to hire into a postdoc position. Postdoc positions are often filled through a passive process where the initial expression of interest from a prospective postdoc is through a “cold-call” contact to a prospective advisor or comes from a recommendation from the PhD advisor or other mentor. An important, but potentially underacknowledged role in this process is provided by the graduate advisor or other mentors of the prospective postdoc, by recommending potential postdoc advisors and/or reaching out to potential advisors.

While undergraduate and graduate admissions often have a centralized application process and faculty searches may have parameters set by a dean or provost’s office, postdoc searches often lack this framework. This lack of systemic structure comes with an accompanying lack of evaluative tools and processes. Centralized processes often require those overseeing the system to decide on what is considered valuable in a candidate and how the candidates will be evaluated. Postdoc hiring, often at the discretion of one faculty member, does not require the same level of decision making or reflection. Additionally, if the faculty member is hiring based on who contacts them at a specific time, they are likely not assessing the full pool of available candidates. This decentralization has far reaching ramifications not only for the hiring process, but also for systematically gathering information about postdoc hiring, since this largely occurs at the level of individual faculty.

The opacity of the processes and practices by which postdocs are identified, recruited, and hired make it difficult to pinpoint where intervention is needed to promote equitable hiring practices and the diversification of the STEM postdoc population. Faculty members can take small steps to help encourage students who might be interested in a postdoc position in their group but may, for a variety of factors, not reach out in a cold email. One way would be a simple statement on a research group encouraging contact from interested potential postdocs as well as instructions for how to make contact. Faculty were asked if they had such a statement on their website. Of the faculty who responded, 42% (11/26; F17) indicated that they have such a statement. Faculty members who had received their PhD more recently were more likely to include a statement on their website with 67% of those who had received their PhD in the past 9 years answering that they did. No faculty member with an academic age 40 or more had a statement. The inclusion of such a statement provides an easy way for the faculty member to signal to prospective postdocs that they are open to hearing from them, and the materials they would like to review. This can help remove some of the mystery from the process and may make the process of sending the initial email less intimidating.

An example of a statement encouraging contact from interested postdocs would be: “Highly motivated postdocs with experience in (specific field) are always encouraged to apply. Please send your CV and a statement of research interests and accomplishments to (email address).” Additional helpful information could include details about what the group looks for in a postdoc, including expertise in a specific field or technique, mentoring of students, etc. A statement signaling that the research group and faculty member prioritize cultivating an inclusive research environment in their group which may include diversity of race/ethnicity, gender identity, neurodivergence, non-traditional educational backgrounds etc. would also be relevant.

Faculty may also want to consider implementing a formal application process for a specific position with information about funding sources and requirements, job expectations, and teaching/mentoring opportunities and advertising it widely. This is done in the UC system, for example, where postdoc applications are collected, and were analyzed by Shauman and Hyunh (2023). Information about funding is especially important. Expectations around funding need to be made very clear at the outset; 42% (27/64; P7) postdocs report having a postdoc advisor who expected them to acquire their own funding for their position, while 38% (10/26; F16) of faculty have hired a postdoc specifically because they had their own funding. If job security is contingent upon acquiring funding, some prospective postdocs may be less willing to take a chance on a job has such an uncertain future, especially if that job involves uprooting their current life, and possibly the lives of their family.

In addition to the efforts of individual faculty, some fields within STEM do widely advertise postdocs through field-specific job boards. Many in math, for example, use mathjobs.com to announce open postdoc opportunities. Astronomy has consolidated faculty and postdoc positions onto the American Astronomical Society Job Register. Outside of field-specific boards, the National Postdoc Association and postdocjobs.com have both created career centers and job boards to advertise available positions in academia, government labs and private industry.

Even if faculty do not have a specific position to advertise, it is still possible to advertise their research group and their desire to hear from interested students. The Research University Alliance recently launched a Postdoc Portal (postdocportal.org) to address some of the concerns raised about postdoc recruiting practices.

When faculty members are recruiting, it is important to keep in mind that campus climate and research group culture are important to postdocs to feel a sense of belonging (Yadav et al, 2020). Faculty may want to consider bringing prospective postdocs to their campus to give them a chance to assess these factors before they commit to a position. They should also be intentional to ensure that postdocs have the opportunity to engage with the larger campus community and to build their own communities and networks. One postdoc surveyed encapsulated all of this when they said “I wished I had gone through a formal interview process (interview, on campus visit, one-on-one discussion with everybody in the group). I also wished I would have spent more time learning the culture of the group and how my skills/background would fit the group dynamic. Making a more standardize[d] way of hiring postdoctoral fellows will help make them feel more appreciated and viewed as true workers and not simply as an extension of their graduate studies.”

This study has examined postdoc hiring practices in STEM within one academic division at one institution. Many areas of inquiry of course remain to be examined. Further expansion to other divisions and institutions would help to understand how various disciplines and institutions may differ from one another. This expansion should include peer institutions across the country as well as institutions who use other models of postdoc hiring. Including a variety of postdoc hiring practices, such as the centralized application system in the UC system and the targeted recruiting of the Stanford Postdoctoral Recruitment Initiative in Science and Medicine (PRISM) will provide the broadest understanding of the current state of postdoc hiring possible. A broader understanding of current postdoc hiring practices is critical to making informed recommendations about best practices for inclusive, diverse hiring that does not discourage or exclude interested parties from the outset. Though many may feel that the way postdoc hiring currently happens works for them, we all have a responsibility to ensure that it works for as many people as possible. Without fully understanding the current practices and their barriers, we cannot begin that process.

## Acknowledgments

We gratefully acknowledge the Research University Alliance Postdoc Working Group for their fruitful discussions around this topic. This material was based upon work supported by the National Science Foundation under Grant Number 2015149. Any opinions, findings, and conclusions or recommendations expressed in this material are those of the author(s) and do not necessarily reflect the views of the National Science Foundation.

Ethics Statement

All participants completed informed consent. This study was reviewed by the Caltech Institutional Review Board (protocol #IR22-1240) and was determined to be exempt from full review.

## Data Availability

The survey questions and responses are provided in Appendices 1 and 2.

## Appendix 1: CCE Survey Questions

**Postdoc Hiring Survey - Postdocs Introduction**

Caltech is a part of the NSF-funded Research University Alliance which, among other goals, seeks to diversify the postdoctoral population at research universities by broadening the networks connecting prospective advisors and postdocs. As part of this process, we need to better understand the current postdoc hiring practices used at our institutions. Though undergraduate, graduate and faculty recruiting processes are well studied in the literature, there is very little available examining the current state of postdoc hiring.

To better understand the mechanisms by which postdocs are hired at Caltech, we are surveying both the faculty and postdocs currently in Division of Chemistry and Chemical Engineering. We ask that you take 5-10 minutes to complete this survey to help us understand how you found your current (and any previous) postdoc position(s). This survey has been reviewed and approved by the Caltech Institutional Review Board. Due to the small size of the population at Caltech, it may be possible for somebody to discover your identity from your responses to certain questions. Your responses will be anonymous and no individually identifying information will be shared in any publications that may result from this survey.

1. Did you complete your undergraduate studies in the United States?

No

Yes

2. Did you complete your PhD in the United States?

No

Yes

3. How many postdoc positions have you held?

1 (this current position)

2

3

4

*Skip To: Q7 If How many postdoc positions have you held? = 1 (this current position)*

4. Were any of your previous postdoc positions in the lab where you completed your PhD?

No

Yes

5. Where did you do your previous postdoc(s)?

6. How did your find your previous postdoc position(s) (excluding position held in PhD lab)? (select all that apply)

Recommendation from PhD advisor/mentor

Previous research based contact

Networking at a conference/meeting

Applied to a posted position

Cold email/phone call to scholar in the field

Applied to a postdoc fellowship

Other (please specify)

7. Have you ever had a postdoc advisor expect you to acquire your own funding for your position?

No

Yes

8. How many years have you been a postdoc at Caltech? less than 1

1

2

3

4

5

6 or more

9. How did you find your current postdoc position?

Recommendation from PhD advisor/mentor

Previous research based contact

Networking at a conference/meeting

Applied to a posted position

Cold email/phone call to scholar in the field

Applied to a postdoc fellowship

Other (please specify)

10. How many positions did you apply to or research groups did you contact when looking for your current position?

11. Did you apply to any specific postdoc position listings?

No

Yes

*Skip To: Q13 If Did you apply to any specific postdoc position listings? = No*

12. Where did you find the position(s) listed?

13. Did your PhD advisor reach out to their network to help you find any of your postdoc positions?

No

Yes

Skip To: Q15 If Did your PhD advisor reach out to their network to help you find any of your postdoc positions? = No

14. Did this result in any potential positions for you?

No

Yes

15. Did you have any personal contact with your current supervisor before you pursued your postdoc opportunity?

No

Yes

16. When did you start looking for your current postdoc position? i.e. about 6 months before your PhD defense, 2 months into your previous 2-year postdoc.

17. Thinking back to your time as a graduate student, was a postdoc position at Caltech something you had pictured for your career path? Why or why not?

18. Have you secured a position for after your postdoc ends? No

Yes

*Skip To: Q20 If Have you secured a position for after your postdoc ends? = No*

19. What is your next position?

Faculty

Another postdoc position

Industry

Additional schooling/training

Non-faculty research position

Something else (please specify)

20. Is there anything else you would like us to know about your experiences in the postdoc hiring process?

21. What is your sex?

Male

Female

Non-binary / third gender

Prefer not to say

**Postdoc Hiring Survey - Faculty Introduction**

Caltech is a part of the NSF-funded Research University Alliance which, among other goals, seeks to diversify the postdoctoral population at research universities by broadening the networks connecting prospective advisors and postdocs. As part of this process, we need to better understand the current postdoc hiring practices used at our institutions. Though undergraduate, graduate and faculty recruiting processes are well studied in the literature, there is very little available examining the current state of postdoc hiring.

To better understand the mechanisms by which postdocs are hired at Caltech, we are surveying both the faculty and postdocs currently in Division of Chemistry and Chemical Engineering. We ask that you take 5-10 minutes to complete this survey to help us understand how you find postdocs in your research group. This survey has been reviewed and approved by the Caltech Institutional Review Board. Due to the small size of the population at Caltech, it may be possible for somebody to discover your identity from your responses to certain questions. Your responses will be anonymous and no individually identifying information will be shared in any publications that may result from this survey.

1. How many years has it been since you earned your PhD?

0 - 9 years (since 2013 or later)

10 - 19 years (2012-2003)

20 - 29 years (2002 - 1993)

30 - 39 years (1992 - 1983)

40+ years (1982 or earlier)

2. How long have you been a faculty member (at any institution)?

0 - 9 years (since 2013 or later)

10 - 19 years (2012 - 2003)

20 - 29 years (2002 - 1993)

30 - 39 years (1992 - 1983)

40+ years (1982 or earlier)

3. Did you hold a postdoc position before you became a faculty member?

No

Yes

*Skip To: Q8 If Did you hold a postdoc position before you became a faculty member? = No*

4. How many postdoc positions have you held?

5. Did you hold a postdoc position at your PhD granting institution?

No

Yes

Skip To: Q6 If Did you hold a postdoc position at your PhD granting institution? = Yes

Skip To: Q7 If Did you hold a postdoc position at your PhD granting institution? = No

6. How did you find your first postdoc position outside of your PhD granting institution? (select all that apply)

Referral from PhD advisor/mentor

Previous research-based contact

Networking at a conference/meeting

Applied to a posted position

Cold email/phone call to scholar in the field

Applied to a postdoc fellowship

Other (please specify)

Skip To: Q8 If Condition: Selected Count Is Greater Than or Equal to 1. Skip To: Have you had, or do you currently have….

7. How did you find your first postdoc position? (select all that apply)

Referral from PhD advisor/mentor

Previous research-based contact

Networking at a conference/meeting

Applied to a posted position

Cold email/call to scholar in the field

Applied to a postdoc fellowship

Other (please specify)

8. Have you had, or do you currently have, postdocs in your research group?

No

Yes

Skip To: End of Survey If Have you had, or do you currently have, postdocs in your research group? = No

9. Approximately how many postdocs have you had in your research group in your faculty career thus far?

0 - 5

6 - 10

11 - 15

16 - 20

21 - 25

26 - 30

31+

10. In the past 15 years, have you ever advertised for a specific postdoc position vacancy in your research group?

No

Yes

11. Which of the following methods do you currently use to find postdocs in your research group? (select all that apply)

Social media (specific position) - LinkedIn, Twitter, Facebook etc.

Social media (general call) - LinkedIn, Twitter, Facebook etc.

Faculty or research group website (specific position)

Faculty or research group website (general call)

Other Caltech website (HR, postdoc office)

Advertising in scientific journal, publication or society

Online job board (i.e. chemistryjobs.com)

Interested postdocs initiate contact (via email, at conferences etc.)

Email announcement of openings to colleagues and research groups with similar interests at other institutions

Recommendation from collaborators or others in the field Other (please specify)

12. In the last 15 years, which of the methods above have resulted in the greatest number of responses/inquiries?

13. In the last 15 years, which of the methods above have resulted in the greatest number of postdoc hires in your lab?

14. In the last 15 years, how many postdocs have you hired where you had no prior personal or professional contact with their PhD or postdoc advisor?

15. If you currently have postdocs in your group, which method(s) did you use to find them? (select all that apply)

Social media (specific position) - LInkedIn, Twitter, Facebook etc.

Social media (general call) - LinkedIn, Twitter, Facebook etc.

Faculty or research group website (specific position)

Faculty or research group website (general call)

Other Caltech website (HR, postdoc office)

Advertising in scientific journal, publication, or society

Online job board (i.e. chemistryjobs.com)

Interested postdocs initiate contact (via email, at conferences, etc.)

Email announcement of openings to colleagues and research groups with similar interests at other institutions

Recommendation from collaborators or others in the field Other (please specify)

16. Have you ever hired a postdoc specifically because they had their own funding?

No

Yes

17. Do you currently have a statement on your website that gives instructions to those interested in joining your group as a postdoc? i.e. Highly motivated postdocs with experience in (specific field) are always encouraged to apply. Please send a CV, 3 of your best publications and a research statement to (email address).

No

Yes

18. Is there anything else you would like us to know about how your find postdocs for your research group?

19. What is your sex?

Male

Female

Non-binary / third gender

Prefer not to say

## Appendix 2: Summary Data Postdoc Hiring Survey - Postdocs

1. Did you complete your undergraduate studies in the United States?

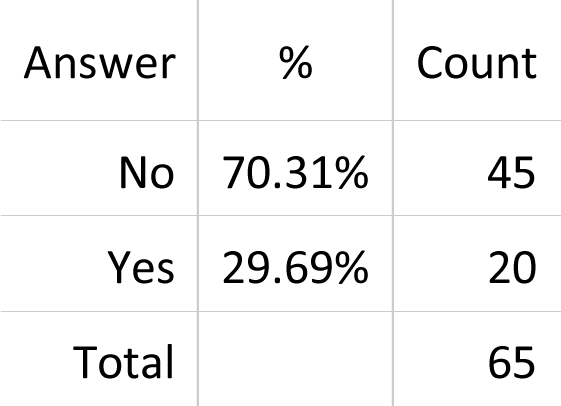

2. Did you complete your PhD in the United States?

| Answer | % | Count |
| --- | --- | --- |
| No | 50.00% | 32 |
| Yes | 50.00% | 33 |
| Total |  | 65 |

3. How many postdoc positions have you held?

| Answer | % | Count |
| --- | --- | --- |
| 1 (this current position) | 75.00% | 49 |
| 2 | 23.44% | 15 |
| 3 | 1.56% | 1 |
| Total |  | 65 |

4. Were any of your previous postdoc positions in the lab where you completed your PhD?

| Answer | % | Count |
| --- | --- | --- |
| No | 43.75% | 7 |
| Yes | 56.25% | 9 |
| Total |  | 16 |

6. How did your find your previous postdoc position(s) (excluding position held in PhD lab)?

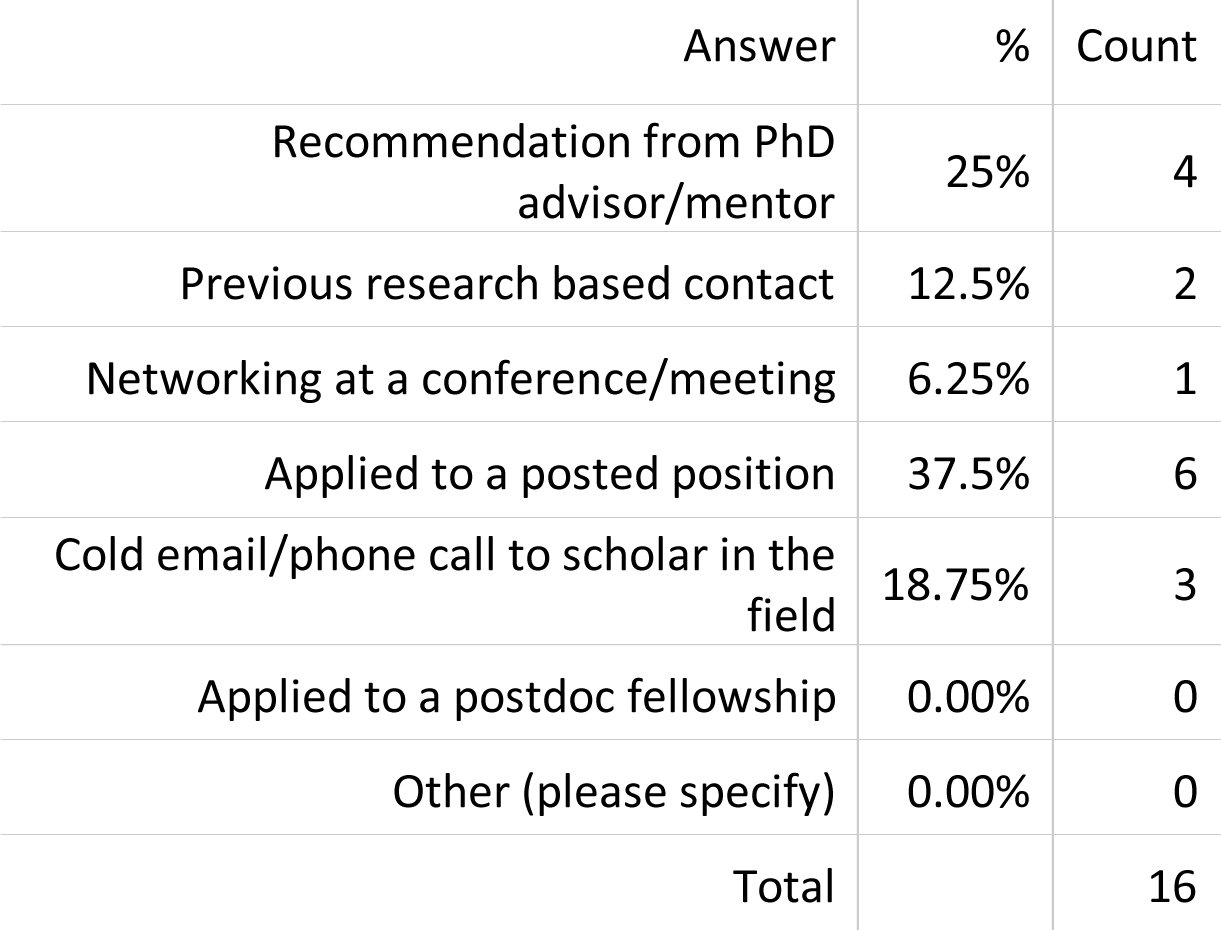

6. How did you find your previous postdoc position(s) (excluding position held in PhD lab)?
| Answer | % | Count |
| --- | --- | --- |
| Recommendation from PhD advisor/mentor | 25% | 4 |
| Previous research based contact | 12.5% | 2 |
| Networking at a conference/meeting | 6.25% | 1 |
| Applied to a posted position | 37.5% | 6 |
| Cold email/phone call to scholar in the field | 18.75% | 3 |
| Applied to a postdoc fellowship | 0.00% | 0 |
| Other (please specify) | 0.00% | 0 |
| Total |  | 16 |

7. Have you ever had a postdoc advisor expect you to acquire your own funding for your position?

| Answer | % | Count |
| --- | --- | --- |
| No | 57.81% | 37 |
| Yes | 42.19% | 27 |
| Total |  | 64 |

8. How many years have you been a postdoc at Caltech?

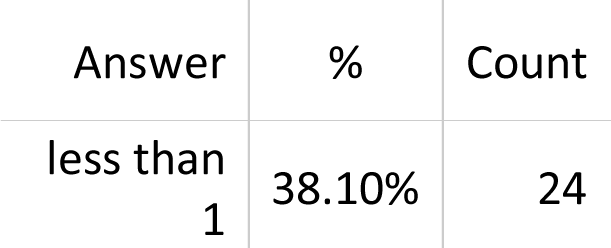
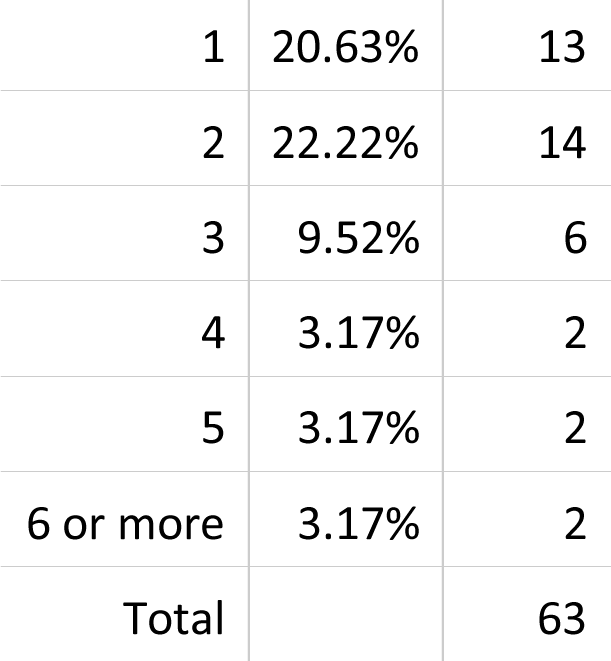

| Answer | % | Count |
| --- | --- | --- |
| less than 1 | 38.10% | 24 |
| 1 | 20.63% | 13 |
| 2 | 22.22% | 14 |
| 3 | 9.52% | 6 |
| 4 | 3.17% | 2 |
| 5 | 3.17% | 2 |
| 6 or more | 3.17% | 2 |
| Total |  | 63 |

9. How did you find your current postdoc position?

| Answer | % | Count |
| --- | --- | --- |
| Recommendation from PhD advisor/mentor | 28.57% | 18 |
| Previous research based contact | 12.70% | 8 |
| Networking at a conference/meeting | 6.35% | 4 |
| Applied to a posted position | 14.29% | 9 |
| Cold email/phone call to scholar in the field | 19.05% | 12 |
| Applied to a postdoc fellowship | 6.35% | 4 |
| Other (please specify) | 12.70% | 8 |
| Total |  | 63 |

10. How many positions did you apply to or research groups did you contact when looking for your current position?
| Answer | % | Count |
| --- | --- | --- |
| 0 | 3.39% | 2 |
| 1 - 5 | 74.58% | 44 |
| 6 - 10 | 6.78% | 4 |
| 11 - 15 | 3.39% | 2 |
| 16 - 20 | 3.39% | 2 |
| 21 - 30 | 3.39% | 2 |
| 31+ | 5.08% | 3 |
| Total |  | 59 |

11. Did you apply to any specific postdoc position listings?

| Answer | % | Count |
| --- | --- | --- |
| No | 82.54% | 52 |
| Yes | 17.46% | 11 |
| Total |  | 63 |

12. Where did you find the position(s) listed?

| Answer (can be more than 1) | Count |
| --- | --- |
| Faculty Website | 3 |
| National Lab Website | 2 |
| Social Media | 2 |
| Job Boards | 2 |
| Email | 1 |
| Word of Mouth | 1 |
| Total | 11 |

13. Did your PhD advisor reach out to their network to help you find any of your postdoc positions?

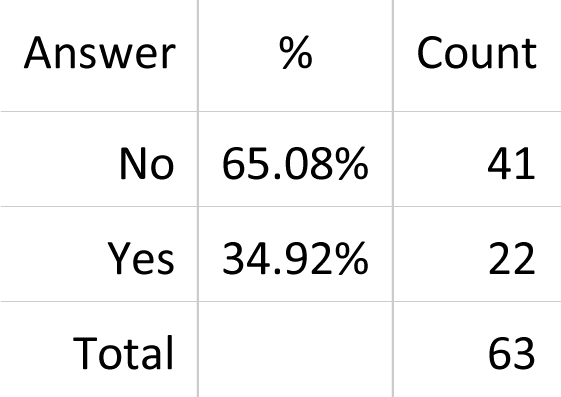

14. Did this result in any potential positions for you?

| Answer | % | Count |
| --- | --- | --- |
| No | 13.64% | 3 |
| Yes | 86.36% | 19 |
| Total |  | 22 |

15. Did you have any personal contact with your current supervisor before you pursued your postdoc opportunity?
| Answer | % | Count |
| --- | --- | --- |
| No | 71.43% | 45 |
| Yes | 28.57% | 18 |
| Total |  | 63 |

| Answer | % | Count |
| --- | --- | --- |
| Before Defense – 0-2 months | 9.09% | 5 |
| Before Defense – 3-5 months | 9.09% | 5 |
| Before Defense – 6-8 months | 32.73% | 18 |
| Before Defense – 9-11 months | 12.73% | 7 |
| Before Defense – 12-18 months | 18.18% | 10 |
| Before End of Postdoc – 0-5 months | 5.45% | 3 |
| Before End of Postdoc – 6-11 months | 5.45% | 3 |
| Before End of Postdoc – 12-18 months | 3.64% | 2 |
| Before End of Postdoc – 19+ months | 3.64% | 2 |
| Total |  | 55 |

18. Have you secured a position for after your postdoc ends?

| Answer | % | Count |
| --- | --- | --- |
| No | 87.30% | 55 |
| Yes | 12.70% | 8 |
| Total |  | 63 |

19. What is your next position?

| Answer | % | Count |
| --- | --- | --- |
| Faculty | 37.50% | 3 |
| Another postdoc position | 12.50% | 1 |
| Industry | 25.00% | 2 |
| Additional schooling/training | 0.00% | 0 |
| Non-faculty research position | 25.00% | 2 |
| Something else (please specify) | 0.00% | 0 |
| Total |  | 8 |

21. What is your sex?

| Answer | % | Count |
| --- | --- | --- |
| Male | 69.84% | 44 |
| Female | 28.57% | 18 |
| Non-binary / third gender | 1.59% | 1 |
| Prefer not to say | 0.00% | 0 |
| Total |  | 63 |

**Postdoc Hiring Survey - Faculty**

1. How many years has it been since you earned your PhD?

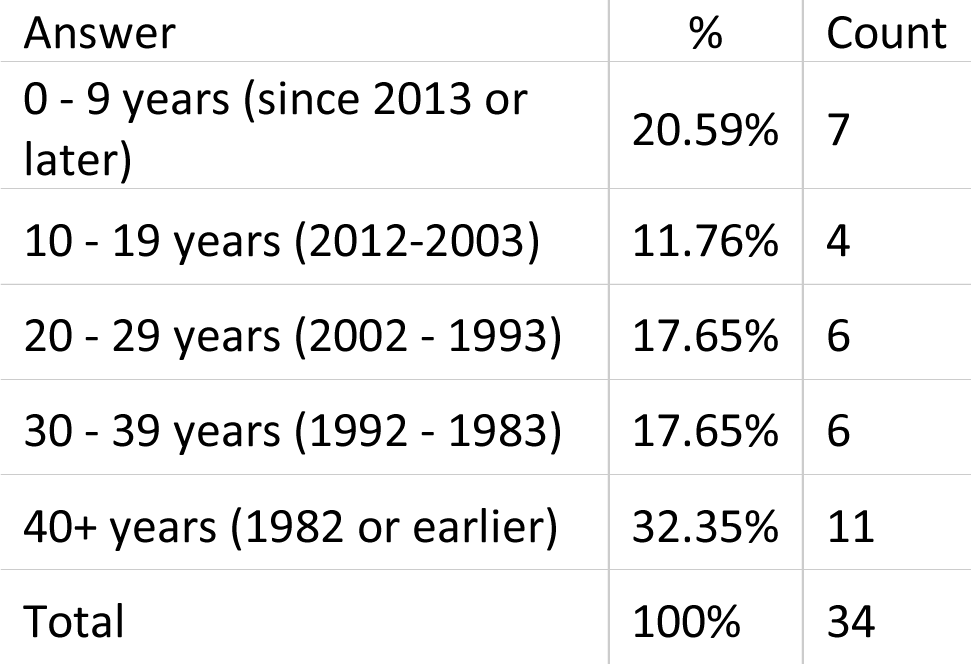

2. How long have you been a faculty member (at any institution)?

| Answer | % | Count |
| --- | --- | --- |
| 0 - 9 years (since 2013 or later) | 23.53% | 8 |
| 10 - 19 years (2012 - 2003) | 14.71% | 5 |
| 20 - 29 years (2002 - 1993) | 14.71% | 5 |
| 30 - 39 years (1992 - 1983) | 23.53% | 8 |
| 40+ years (1982 or earlier) | 23.53% | 8 |
| Total |  | 34 |

3. Did you hold a postdoc position before you became a faculty member?

| Answer | % | Count |
| --- | --- | --- |
| No | 17.65% | 6 |
| Yes | 82.35% | 28 |
| Total |  | 34 |

4. How many postdoc positions have you held?

| Answer | % | Count |
| --- | --- | --- |
| 1 | 75.00% | 21 |
| 2 | 25.00% | 7 |
| Total |  | 28 |

5. Did you hold a postdoc position at your PhD granting institution?

| Answer | % | Count |
| --- | --- | --- |
| No | 75.00% | 21 |
| Yes | 25.00% | 7 |
| Total |  | 28 |

6&7. How did you find your first postdoc position (outside of your PhD granting institution)?

| Answer (can choose more than 1) | Count |
| --- | --- |
| Referral from PhD advisor/mentor | 8 |
| Previous research-based contact | 2 |
| Networking at a conference/meeting | 1 |
| Applied to a posted position | 0 |
| Cold email/phone call to scholar in the field | 13 |
| Applied to a postdoc fellowship | 5 |
| Other (please specify) | 3 |
| Total | 32 |

8. Have you had, or do you currently have, postdocs in your research group?

| Answer | % | Count |
| --- | --- | --- |
| No | 7.14% | 2 |
| Yes | 92.86% | 26 |
| Total |  | 28 |

9. Approximately how many postdocs have you had in your research group in your faculty career thus far?
| Answer | % | Count |
| --- | --- | --- |
| 0 - 5 | 26.92% | 7 |
| 6 - 10 | 7.69% | 2 |
| 11 - 15 | 15.38% | 4 |
| 16 - 20 | 11.54% | 3 |
| 21 - 25 | 7.69% | 2 |
| 26 - 30 | 7.69% | 2 |
| 31+ | 23.08% | 6 |
| Total |  | 26 |

10. In the past 15 years, have you ever advertised for a specific postdoc position vacancy in your research group?
| Answer | % | Count |
| --- | --- | --- |
| No | 38.46% | 10 |
| Yes | 61.54% | 16 |
| Total |  | 26 |

11. Which of the following methods do you currently use to find postdocs in your research group?

| Answer (can chose more than 1) | Count |
| --- | --- |
| Social media (specific position) - LinkedIn, Twitter, Facebook etc. | 7 |
| Social media (general call) - LinkedIn, Twitter, Facebook etc. | 5 |
| Faculty or research group website (specific position) | 6 |
| Faculty or research group website (general call) | 9 |
| Other Caltech website (HR, postdoc office) | 1 |
| Advertising in scientific journal, publication or society | 4 |
| Online job board (i.e. chemistryjobs.com) | 2 |
| Interested postdocs initiate contact (via email, at conferences etc.) | 21 |
| Email announcement of openings to colleagues and research groups with similar interests at other institutions | 10 |
| Recommendation from collaborators or others in the field | 15 |
| Other (please specify) | 2 |
| Total | 82 |

12. In the last 15 years, which of the methods above have resulted in the greatest number of responses/inquiries?
| Answer (can chose more than 1) | Count |
| --- | --- |
| Social media - LinkedIn, Twitter, Facebook etc. | 4 |
| Faculty or research group website | 1 |
| Other Caltech website (HR, postdoc office) | 0 |
| Advertising in scientific journal, publication or society | 1 |
| Online job board (i.e. chemistryjobs.com) | 2 |
| Interested postdocs initiate contact (via email, at conferences etc.) | 16 |
| Email announcement of openings to colleagues and research groups with similar interests at other institutions | 1 |
| Recommendation from collaborators or others in the field | 2 |
| Word of Mouth | 1 |
| Total | 28 |

13. In the last 15 years, which of the methods above have resulted in the greatest number of postdoc hires in your lab?
| Answer (can chose more than 1) | Count |
| --- | --- |
| Social media - LinkedIn, Twitter, Facebook etc. | 1 |
| Faculty or research group website | 0 |
| Other Caltech website (HR, postdoc office) | 0 |
| Advertising in scientific journal, publication or society | 1 |
| Online job board (i.e. chemistryjobs.com) | 1 |
| Interested postdocs initiate contact (via email, at conferences etc.) | 18 |
| Email announcement of openings to colleagues and research groups with similar interests at other institutions | 0 |
| Recommendation from collaborators or others in the field | 4 |
| Word of Mouth | 1 |
| Total | 26 |

| Answer | Count |
| --- | --- |
| 0 - 5 | 17 |
| 6 | 1 |
| 15 | 1 |
| 20+ | 3 |
| Less than half | 1 |
| Most of them | 2 |
| Total | 25 |

15. If you currently have postdocs in your group, which method(s) did you use to find them?

| Answer | Count |
| --- | --- |
| Social media (specific position) - LinkedIn, Twitter, Facebook etc. | 4 |
| Social media (general call) - LinkedIn, Twitter, Facebook etc. | 0 |
| Faculty or research group website (specific position) | 2 |
| Faculty or research group website (general call) | 2 |
| Other Caltech website (HR, postdoc office) | 0 |
| Advertising in scientific journal, publication, or society | 2 |
| Online job board (i.e. chemistryjobs.com) | 1 |
| Interested postdocs initiate contact (via email, at conferences, etc.) | 19 |
| Email announcement of openings to colleagues and research groups with similar interests at other institutions | 3 |
| Recommendation from collaborators or others in the field | 8 |
| Other (please specify) | 3 |
| Total | 44 |

16. Have you ever hired a postdoc specifically because they had their own funding?

| Answer | % | Count |
| --- | --- | --- |
| No | 61.54% | 16 |
| Yes | 38.46% | 10 |
| Total | 100% | 26 |

| Answer | % | Count |
| --- | --- | --- |
| No | 57.69% | 15 |
| Yes | 42.31% | 11 |
| Total |  | 26 |

19. What is your sex?

| Answer | % | Count |
| --- | --- | --- |
| Male | 69.23% | 18 |
| Female | 26.92% | 7 |
| Non-binary / third gender | 0.00% | 0 |
| Prefer not to say | 3.85% | 1 |
| Total |  | 26 |

## References

Andalib, M. A., N. Ghaffarzadegan and R. C. Larson (2018). “The Postdoc Queue: A Labour Force in Waiting.” Syst. Res. 35: 675–686.

Denton M., M. Borrego and D.B. Knight (2022) “U.S. postdoctoral careers in life sciences, physical sciences and engineering: Government, industry, and academia”. PLoS ONE 17: e0263185

Eaton, A. A., J. F. Saunders, R. K. Jacobson and K. West (2020). “How Gender and Race Stereotypes Impact the Advancement of Scholars in STEM: Professors’ Biased Evaluations of Physics and Biology Post-Doctoral Candidates.” Sex Roles 82(3): 127–141.

Grinstein, A. and R. Treister (2018). “The unhappy postdoc: a survey based study [version 2; peer review: 2 approved, 1 approved with reservations, 1 not approved].” F1000Research 6: 1642.

Huynh, J. and K. A. Shauman. (2021). “Postdoctoral Hiring & Equity Issues in STEM: Employment Trends, Policy, and Research. Inclusive Graduate Education Network Research Brief” from equitygraded.org/wp-content/uploads/2022/04/Hiring-Brief.pdf.

König, J. (2022). “Postdoctoral employment and future non-academic career prospects.” PLoS ONE 17: e0278091.

Lambert, W. M., M. T. Wells, M. F. Cipriano, J. N. Sneva, J. A. Morris and L. M. Golightly (2020). “Career choices of underrepresented and female postdocs in the biomedical sciences.” eLife 9: e48774.

Langin, K. (2022). “U.S. labs face severe postdoc shortage” Science 376: 1369–1370.

Liera, R and C. Ching (2020). “Reconceptualizing “merit” and “fit”: An equity-minded approach to hiring”. In A. Kezar & J. Posselt (Eds.) Higher education administration for social justice and equity: Critical perspectives for leadership (pp. 11–131). New York, NY: Routledge.

Matias, J.N., N.A. Lewis and E.C. Hope (2022). “US universities are not succeeding in diversifying faculty” Nature Human Behav. 6: 1606–1608.

MIT Postdoctoral Association, Diversity, Equity, and Inclusion Committee (2020). “Postdoc Hiring and Recruitment at MIT” downloaded from https://drive.google.com/file/d/1bhQqdNUzSbzn3t8K-BREWMhkr3Xn5W_9/view

National Academies of Sciences, Engineering and Medicine (2023). Advancing Antiracism, Diversity, Equity and Inclusion in STEMM Organizations: Beyond Broadening Participation. Washington, DC, The National Academies Press.

National Academy of Sciences, National Academy of Engineering, and Institute of Medicine (2000). Enhancing the Postdoctoral Experience for Scientists and Engineers: A Guide for Postdoctoral Scholars, Advisers, Institutions, Funding Organizations, and Disciplinary Societies. Washington, DC, The National Academies Press.

National Academy of Sciences, National Academy of Engineering, and Institute of Medicine (2014). The Postdoctoral Experience Revisited. Washington, DC, The National Academies Press.

National Association of College Admissions Counseling (2023). “Elevate equity 2023: Summary and framework. Available: https://www.nacacnet.org/wp-content/uploads/ElevateEquityReport.pdf.

Patt, C., Eppig, A. and Richards, M.A. (2022). “Postdocs as key to faculty diversity: A structured and collaborative approach for research universities.” Frontiers in Psychology 12, article 759263, doi: 10.3389/fpsyg.2021.759263.

Posselt, J. R. (2016). “Inside graduate admissions: Merit, diversity, and faculty gatekeeping”. Cambridge, MA: Harvard University Press.

Posselt, J.R. (2020). “Equity in science: Representation, culture and the dynamics of change in graduate education”. Stanford, CA: Stanford University Press.

Sauermann, H. and M. Roach (2016). “Why pursue the postdoc path?” Science 352(6286): 663–664.

Sensoy, Ö. And R. DiAngelo (2017). “”We are all for diversity, but…”: How faculty hiring committees reproduce whiteness and practical suggestions for how they can change.” Harvard Educational Review, 87(4):557–580.

Shauman, K. A. and J. Huynh (2023). “Gender, race-ethnicity and postdoctoral hiring in STEMM fields.” Social Science Research 113: 102854.

Yadav, A., C.D. Seals, C.M. Soto Sullivan, M. Lachney, Q. Clark, K.G. Dixon, and M.J.T. Smith (2020) “The forgotten scholar: Underrepresented minority postdoc experiences in STEM fields” Educ. Studies 56: 160–185.

